# Divergent viral phosphodiesterases for immune signaling evasion

**DOI:** 10.1101/2025.08.21.671373

**Authors:** Erin E. Doherty, Jason Nomburg, Benjamin A. Adler, Santiago Lopez, Kendall Hsieh, Nathan Price, Nurashau Blount, Jennifer A. Doudna

## Abstract

Cyclic dinucleotides (CDNs) and other short oligonucleotides play fundamental roles in immune system activation in organisms ranging from bacteria to humans. In response, viruses use phosphodiesterase-mediated oligonucleotide cleavage for immune evasion, a strategy whose diversity has not yet been explored. We used a canonical 2H phosphodiesterase (2H PDE) structure-based search of prokaryotic and eukaryotic viral sequences to identify an exceptional diversity of 2H PDEs across the virome, including enzymes not detectable with sequence search methods alone. Despite active site conservation, biochemical experiments revealed remarkable substrate specificity of these PDEs that corresponds to variation in the core 2H fold. This nuanced specificity allows 2H PDEs to selectively degrade oligonucleotide messengers to avoid interfering with host immune signaling. Together, these findings nominate viral 2H PDEs as key regulators of CDN signaling across the tree of life.

## Introduction

cGAS/DncV-like nucleotidyltransferases (CD-NTases) are a critical component of the innate immune response in humans and bacteria^1,2^. These enzymes produce nucleotide messengers in response to viral cues, which in turn activate an array of antiviral effector proteins. Humans encode two cGAS-like enzymes: cGAS, which responds to cytoplasmic double-stranded DNA to produce 2’3’ cyclic GMP-AMP (cGAMP); and oligo-adenylate synthase (OAS) proteins, which respond to double-stranded RNA to produce linear oligoadenylates with 2’5’ linkages (2’5’ OA) ^3^. Bacteria and archaea also encode cGAS-like enzymes, where they sense viral infection as part of cyclic oligonucleotide-based antiphage signaling system (CBASS) defense systems ^4^.

Viruses use diverse strategies to evade CD-NTases and facilitate viral replication. These strategies include directly targeting the CD-NTase proteins themselves, degrading or sequestering nucleotide messengers produced by CD-NTases, or targeting downstream effector proteins. Degradation of nucleotide messengers by phosphodiesterases (PDEs) is a notable mechanism through which viruses evade CD-NTase sensing. This includes the protein anti-cbass 1 (Acb1), a phage PDE that cleaves cGAMP and can promote phage infection in the context of CBASS sensing ^5^. Acb1 contains a two histidine (2H) phosphodiesterase fold, present in diverse cellular enzymes within and beyond humans ^6^. We recently found that eukaryotic viruses such as avian poxviruses and iridoviruses also encode 2H PDEs that cleave cGAMP, suggesting that cGAMP degradation by 2H PDEs is a pan-viral mechanism of immune evasion ^7,8^.

CD-NTases in bacterial CBASS systems produce diverse nucleotide messengers beyond cGAMP ^4,9^. Bacteria contain hundreds of other viral sensing pathways, including many that involve nucleotide messengers ^10^. Furthermore, many signaling pathways involved in cellular homeostasis in both eukaryotes and prokaryotes also involve nucleotide messengers ^11^. However, whether and how viral phosphodiesterases interact with these diverse nucleotide signals is not clear. Furthermore, the possibility of diverse phage-encoded 2H PDEs beyond Acb1 has not been explored. Here, we used structure-guided database searches to identify dozens of novel 2H PDE clades, revealing enzymes with a range of activities against diverse nucleotide messengers involved in cellular homeostasis and immunity. These results suggest that viruses have adapted the 2H PDE core structure repeatedly to enable evasion of diverse bacterial immune systems, making 2H PDEs key players in the virus-host conflict across the tree of life.

## Results

### Structure-guided discovery of viral 2H PDEs

CD-NTases sense viral cues and produce diverse nucleotide messengers to stimulate an anti-viral response ^1,2^ (Fig. 1A). We previously found that 2H PDEs are used by some animal and bacterial viruses to degrade nucleotide messengers produced by CD-NTases ^7,8^ (Fig. 1B). This raises the possibility that additional viruses encode 2H PDEs involved in evasion of CD-NTase immunity. To address this hypothesis, we used the protein structure search algorithm DALI ^12^ to align the predicted structures of eight known viral 2H PDEs against a collection of predicted structures of phage proteins. This led to 84 alignments, including many aligned proteins with less than 20% sequence identity but strong structural similarity (marked by DALI Z scores above 5) to a viral 2H (Fig. 1C). Many of these alignments fall below the “twilight zone” of sequence similarity, past which it is challenging to find true protein similarity ^13^. Structural alignments provide a substantial improvement over a similar search with Jackhmmer^14^, a sensitive sequence-based aligner (Supplementary Fig. 1A), showing the benefits of protein structure search over sequence-based methods. Of these 84 alignments, manual curation revealed 75 hits with intact catalytic histidines and a conserved 2H core fold (Supplementary Table 1). Together, this analysis revealed substantial numbers of phage 2H PDEs that have not been studied or even identified previously.

**Figure 1.**
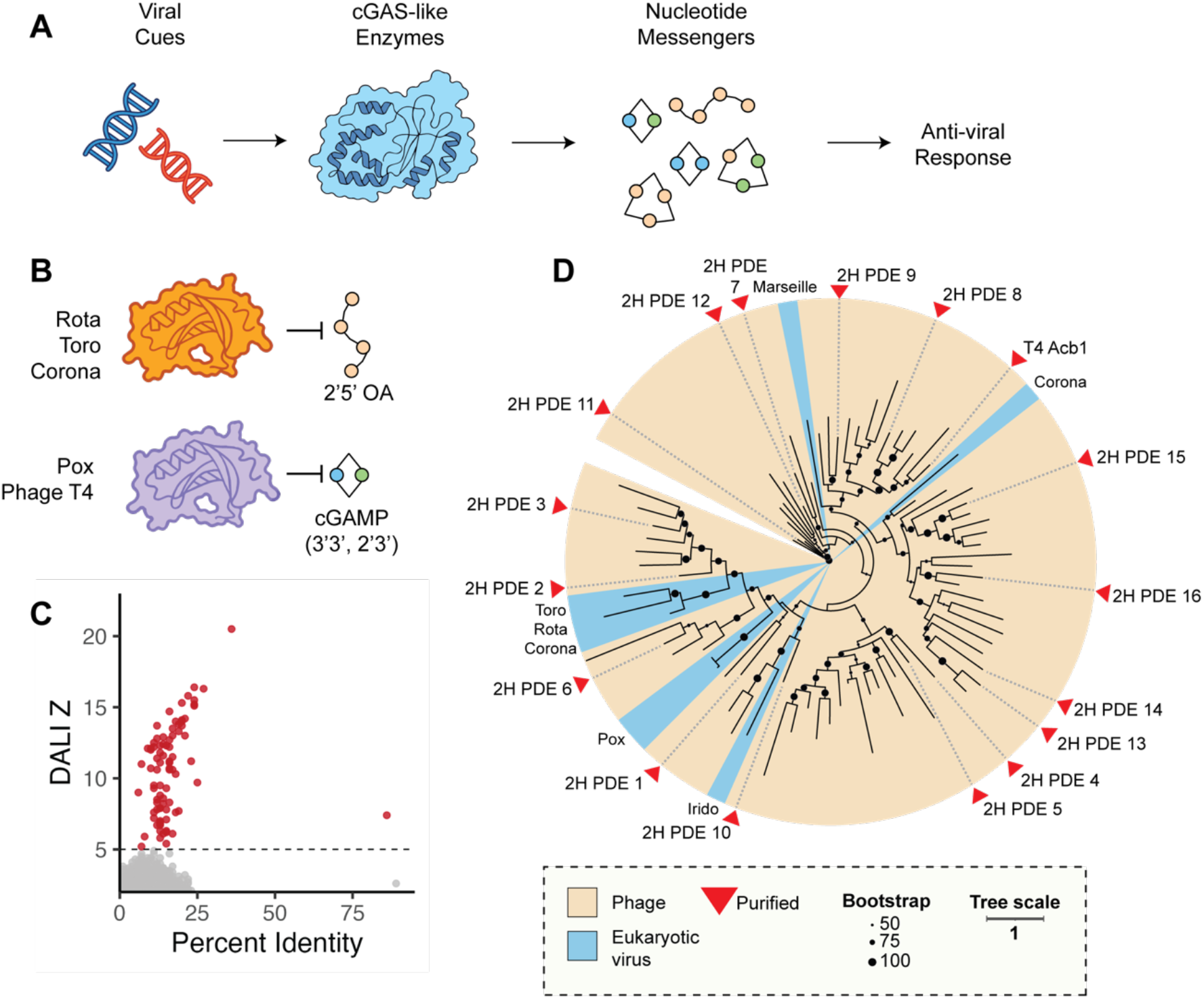
Protein structure search reveals the diversity of 2H PDEs across the virome. **A**. CD-NTases are activated by a viral cue, often DNA or RNA. Upon recognition, they synthesize nucleotide messengers responsible for downstream signalling. **B**. 2H PDEs encoded by eukaryotic RNA viruses degrade 2’5’OA, produced by OAS-like CD-NTases. Avian poxvirus 2H PDEs and enzymes such as phage T4 Acb1 (noted in red text) can degrade cGAMP. **C**. Protein structure search using DALI reveals many phage proteins with structural similarity to known viral 2H PDEs. The X axis indicates percentage sequence identity, while the Y axis indicates the DALI Z score. Each dot is a target protein, and proteins with a Z score higher than 5 are colored red. The “twilight zone” of protein sequence similarity, below approximately 30% sequence similarity, is marked in blue. **D**. A phylogenetic tree that incorporates 2H PDEs found in eukaryotic viruses and phage. The highlighted color indicates whether the PDE is found in phage or eukaryotic viruses, and bootstrap values 50 or higher are indicated by dots on each branch. Proteins for which we attempted downstream purification are indicated with red triangles.

Next, we investigated the evolutionary relationships of these 2H PDEs with known 2H lineages. Low sequence similarity can make it challenging to generate accurate multi-sequence alignments (MSAs) using conventional sequence-based methods. To address this challenge, Foldmason ^15^ was applied to use protein structure alignments to guide MSA generation of viral 2H PDEs and used this MSA to generate a phylogenetic tree using IQ-Tree ^16^ (Fig. 1D). This revealed that viral 2H PDEs are extremely diverse, and include many phage 2H PDE lineages that are distinct from T4 Acb1. Of the 75 detected 2H PDEs, 13 reside in proteins that contain one or more putative additional domains, including the haloacid dehydrogenase superfamily of enzymes, ribonucleases, acetyltransferases, hydrolases, proteases, and domains of unknown function (Supplementary Table 2).

### Viral 2H PDEs cleave diverse nucleotide substrates

To investigate the structural diversity of 2H PDEs, we conducted all-by-all structural alignments with DALI. This revealed that viral 2H PDEs are structurally heterogenous, and fall within distinct structural groups (Fig. 2A,B). We reasoned that this structural variation may enable diversification of biochemical activity and cleavage of diverse nucleotide substrates. Oligonucleotide messengers known to function in immunity differ in the identity of the nucleotide base(s), linkage chemistry, and whether the molecule is cyclized or linear ^10,17^. To determine the breadth of targeting by viral 2H PDEs, we carried out in vitro assays that assessed their ability to target oligonucleotide-based messengers from different immune systems.

**Figure 2.**
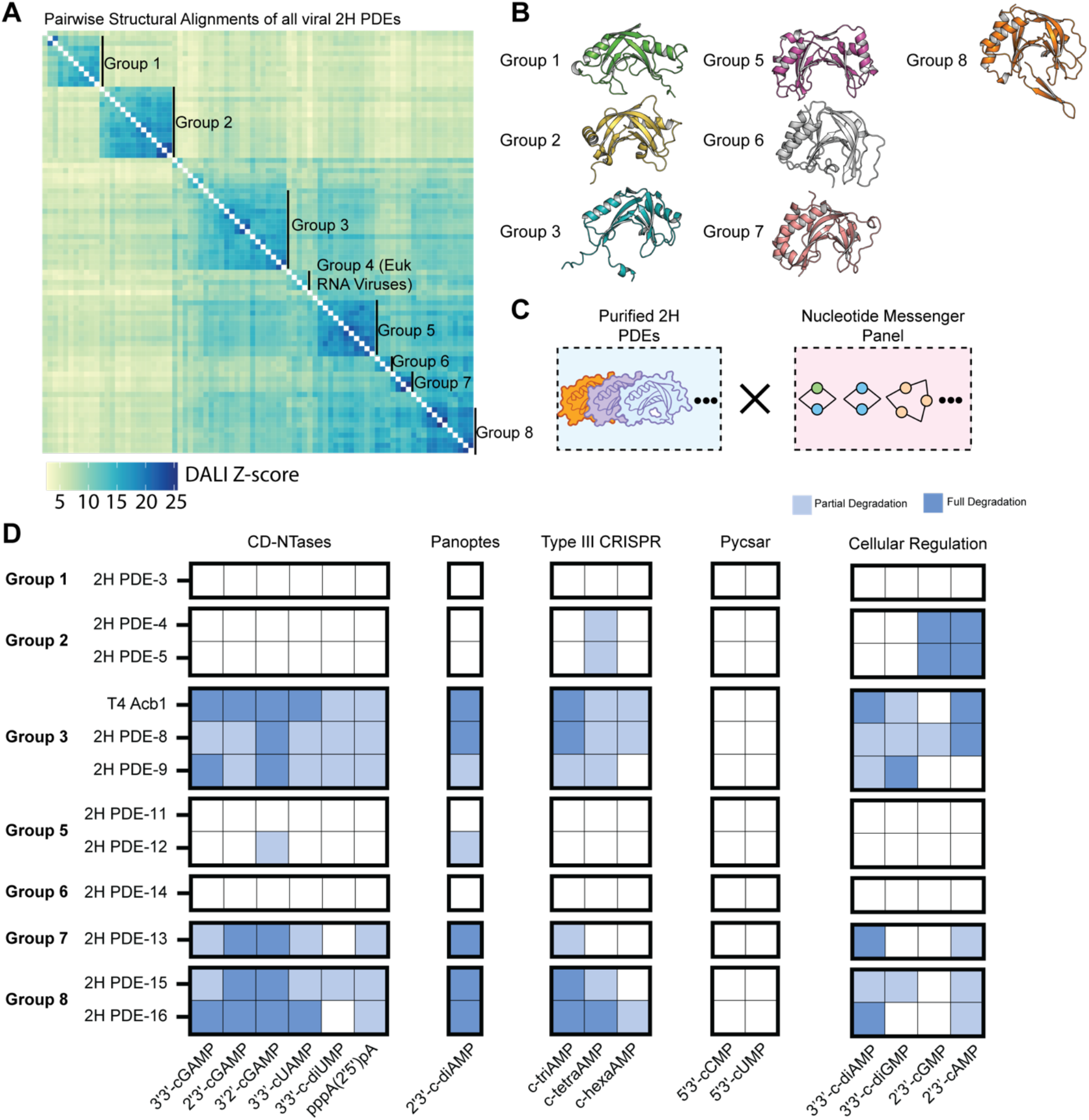
Structural variation in the 2H core is associated with biochemical specificity. **A**. All-by-all alignments between all viral 2H PDEs using DALI, with the color indicating the DALI Z score. Clusters of 2H PDEs with high pairwise Z scores were manually classified into structure groups. **B**. Representative 2H PDE structures from each structure group. **C**. Representative 2H PDEs from some structure groups were purified, and their ability to cleave a panel of 18 nucleotide messengers was assessed. **D**. The ability of 12 viral 2H PDEs to degrade 18 different nucleotide messengers was assessed, with full degradation indicated in dark blue, partial degradation indicated in light blue, and no observable degradation indicated by a white square. Degradation was read out through thin-layer chromatography in technical duplicate. Nucleotide messengers are indicated on the X axis, and include those produced during cellular regulation as well as the bacterial defense systems involving CD-NTases, Panoptes, Type III CRISPR, and Pycsar.

We purified 16 candidate 2H PDEs (Fig. 1D) representing seven distinct structural groups (Fig. 2A, B). Each PDE was tested against a panel of oligonucleotide messengers involved in different signaling pathways: CD-NTase-related molecules (cGAMP isomers, cUAMP, c-diUMP, and 2’5’ linked diadenylate) ^10^, Type III CRISPR (cyclic tri-, tetraand hexaadenylate) ^18,19^, Panoptes (2’3’-c-diAMP) ^20,21^, Pycsar (5’3’-cCMP/UMP) ^22^, and homeostatic regulation (3’3’-c-diAMP/GMP, and 2’3’-c-AMP/GMP) ^23–25^. PDEs were evaluated for activity against this panel by using thin layer chromatography (TLC) to assay for degradation of the messenger molecules after a 5-minute incubation (Fig. 2C, Supplementary Fig. 2). Strikingly, the different viral 2H PDEs displayed unique substrate scope and preferences that correlated with structural groups (Fig. 2D, Supplementary Fig. 3,4). Some structural groups (group 1 and group 6) did not display activity against any molecules in our assay conditions, suggesting that their natural substrate(s) may not be included in this diverse panel of phosphodiesterase-linked chemical messengers. Structural groups 3 (containing T4 Acb1), 7, and 8 showed broad spectrum activity against molecules involved in CD-NTase pathways, cellular regulation and Type III CRISPR systems. However, there were also groups that were narrowly confined to specific messenger molecules (group 2 and group 5), suggesting that restricting activity to select targets may offer an advantage in certain cases.

**Figure 3.**
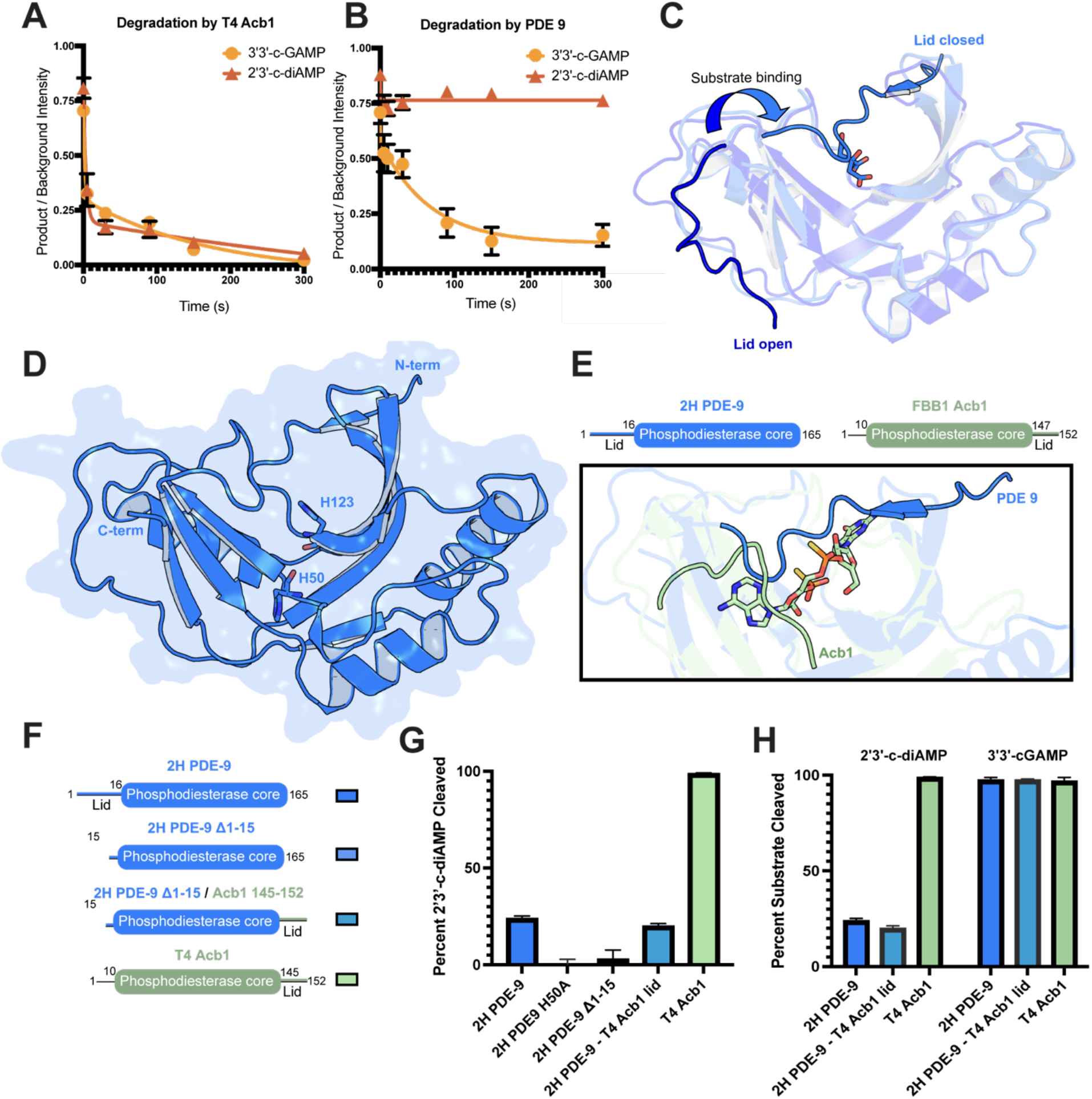
The 2H PDE enzyme core determines differential substrate specificity. **A**. Degradation of 3’3’-cGAMP (light orange) and 2’3’-c-diAMP (dark orange) by T4 Acb1. The plot represents the percent of substrate remaining over time (in seconds) compared to a chemical standard as determined by TLC. Values plotted are the mean of three technical replicates and error bars represent the standard deviation. **B**. Degradation of 3’3’cGAMP (light orange) and 2’3’-c-diAMP (dark orange) by 2H PDE-9. The plot represents the percent of substrate remaining over time (in seconds) compared to a chemical standard as determined by TLC. Values plotted are the mean of three technical replicates and error bars represent the standard deviation. **C**. Overlay of 2H PDE-9 structures in the apo / open loop form (PDE-9 H50A) (dark blue, PDB ID: 9Q2Y, this work) and citrate bound / closed loop form (light blue, PDB ID: 9Q2G, this work). Citrate is shown in the active site in light blue and red. The arrow indicates the change in the lid domain positioning upon substrate binding. Lid open/closed labels are adjacent to the N-terminal residues of each structure. **D**. Structure of 2H PDE-9 in the closed loop form (PDB ID: 9Q2G) with N and C terminus indicated. Catalytic histidines (H50 and H123) are shown in the active site. E. Domain organization of 2H PDE-9 (blue) and FBB1 Acb1 (green) indicating the positioning of the lid domain on each enzyme (PDE-9, N-terminal; Acb1, C-terminal). **F**. Domain organization of 2H-PDE mutants (wild-type, Δ1-15, Δ1-15 + Acb1 147-152) and T4 Acb1. **G**. Cleavage of 2’3’-c-di-AMP over time by enzymes from (**Fig. 3F**). Values plotted are the percent of substrate depleted by each PDE as determined by TLC. Values plotted are the mean of three technical replicates and error bars represent the standard deviation. **H**. Cleavage of 3’3’-cGAMP over time by enzymes from (**Fig. 3F**). Values plotted are the percent of substrate depleted by each PDE as determined by TLC. Values plotted are the mean of three technical replicates and error bars represent the standard deviation.

**Figure 4.**
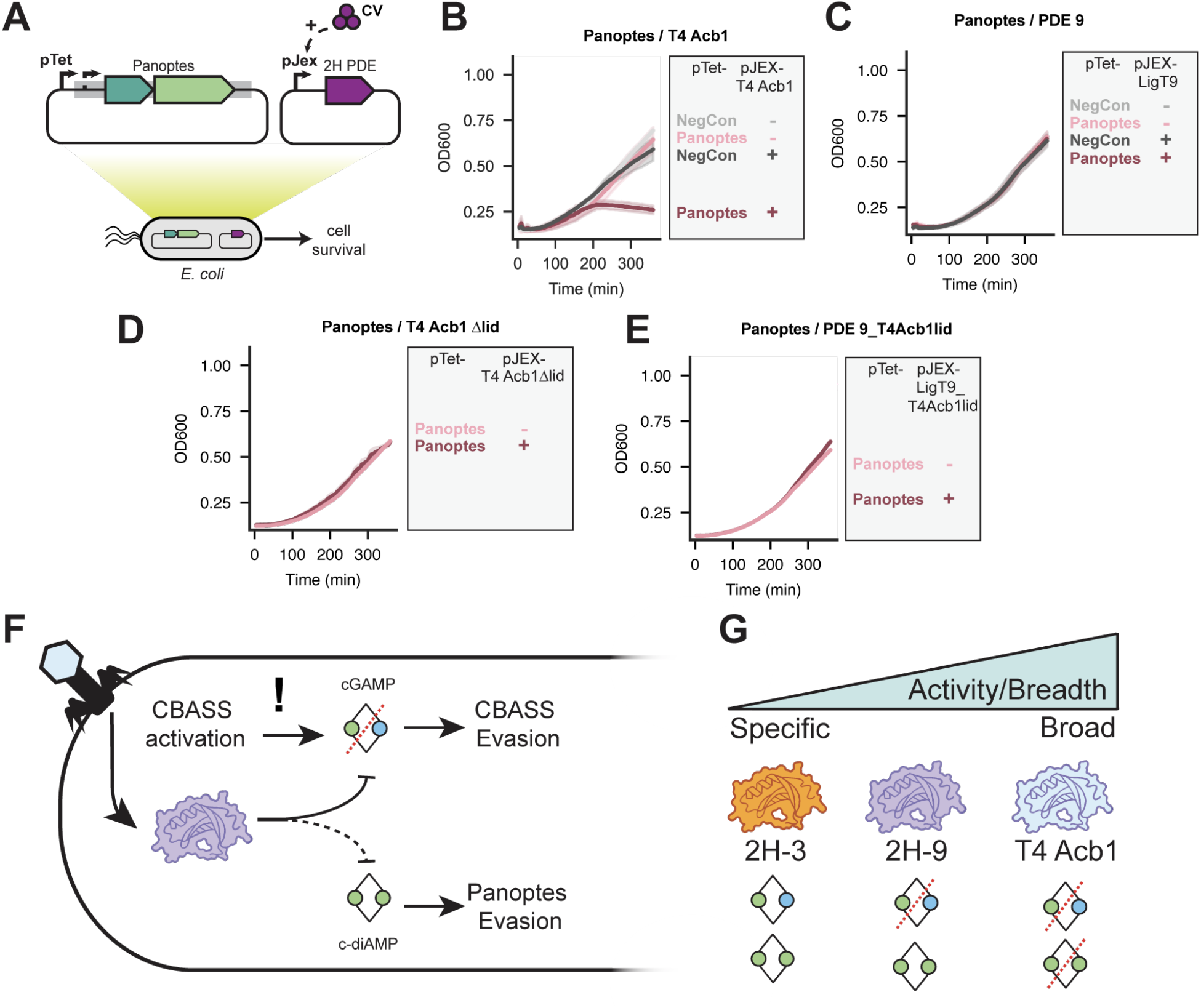
The 2H PDE enzyme core determines differential substrate specificity. **A**. Panoptes activator assays in *E. coli* to determine cell survival. The Panoptes system (mCpol, dark green; 2TMβ, light green) is under a pTet promoter in the context of its native promoter and candidate activators (2H PDEs) are under a pJEX inducible promoter subject to crystal violet (CV) induction). **B**. Panoptes activator assay where the candidate activator, T4 Acb1, is under a pJEX inducible promoter. Under the pTet promoter, NegCon indicates a vector control plasmid and Panoptes indicates the same plasmid containing the Panoptes system in its native context. Conditions involved no induction of the pJEX promoter (−) or the addition of 250 nM crystal violet (+). Graphs plot the cell density (OD600) over time (in minutes). The solid line indicates the mean of three biological replicates and the shaded region represents the standard deviation. **C**. Panoptes activator assay where the candidate activator, T4 Acb1, is under a pJEX inducible promoter. Under the pTet promoter, NegCon indicates a vector control plasmid and Panoptes indicates the same plasmid containing the Panoptes system in its native context. Conditions involved no induction of the pJEX promoter (−) or the addition of 250 nM crystal violet (+). Graphs plot the cell density (OD600) over time (in minutes). The solid line indicates the mean of three biological replicates and the shaded region indicates the standard deviation. **D**. Panoptes activator assay with T4 Acb1 Δ145-152 (deletion of lid residues or Δlid) under the pJEX promoter in the Panoptes activator assay. Panoptes is under the Tet promoter in its native context, and a plasmid with the pJEX promoter contains T4 Acb1 Δ145-152. Conditions involved no induction of the pJEX promoter (−) or the addition of 250 nM crystal violet (+). Graphs plot the cell density (OD600) over time (in minutes). The solid line indicates the mean of three biological replicates and the shaded region indicates the standard deviation. **E**. PDE-9 Δ1-15 + T4 Acb1 145-152 (PDE-9_T4Acb1lid) (**Fig. 3F**) under the pJEX promoter in the Panoptes activator assay. Panoptes is expressed under the pTet promoter in its native context. Conditions involved no induction of the pJEX promoter (−) or the addition of 250 nM crystal violet (+). Graphs plot the cell density (OD600) over time (in minutes). The solid line indicates the mean of three biological replicates and the shaded region indicates the standard deviation. **F**. Phage infection triggers activation of CBASS to produce cyclic oligonucleotides (3’3’-cGAMP). Cleavage of 3’3’-cGAMP by 2H PDE-9 leads to CBASS evasion. Avoidance of 2’3’-c-diAMP by 2H PDE-9 avoids activation of the Panoptes system. **G**. Specificity vs. breadth of cyclic oligonucleotide targeting by 2H PDEs.

Variation in the migration patterns of post-reaction products reflected differences in the nature of degradation across PDEs (Supplementary Fig. 5). Analysis of the molecular composition of 3’3’-cGAMP reaction products revealed that cleavage by different PDEs produced distinct product profiles (Supplementary Fig. 6). Although a conserved intermediate (GpA>p, Supplementary Fig. 7) was detected in each reaction, differences in subsequent product identity may influence the extent of immune suppression, persistence of signaling intermediates or host recognition of degradation products – providing a nuanced layer of specificity in phage anti-defense. Product distributions were not fully conserved even within structural groups (e.g. structural group 3: 2H PDE-8, 9, and T4 Acb1) suggesting that small differences in active site architecture or substrate positioning have functional consequences on oligonucleotide recognition.

### PDE core governs substrate selectivity

Structural group 3 (T4 Acb1, 2H PDE-8, and 2H PDE-9) exhibited the broadest substrate tolerance when tested against the nucleotide messenger panel (Fig. 2D). T4 Acb1 and 2H PDE-9 both efficiently degrade 3’3’-cGAMP – the molecule produced by a large fraction of CBASS systems ^9^. However, there were notable differences between the ability of 2H PDE-9 and T4 Acb1 to target 2’3’-c-di-AMP (Supplementary Fig. 3). To explore the extent of this difference in substrate preference, we performed a TLC-based kinetic assay to observe the rate of degradation by treating 1.25 mM substrate with 8 µM PDE (Supplementary Fig. 8). While Acb1 completely consumed 2’3’-c-di-AMP (k_obs_ = 0.23 ± 0.06 min^-1^) over 5 minutes (Fig. 3A), 2H PDE-9 reaches less than 25% conversion over the same time period (Fig. 3B). To determine if the difference in rate was substrate specific, or an intrinsic quality of the enzyme, we also determined the rate of 3’3’-cGAMP degradation by both enzymes. Acb1 and 2H PDE-9 both exhibited robust activity against 3’3’-cGAMP (k_obs_ = 0.21 ± 0.12 and 0.02 ± 0.01 min^-1^ respectively). Acb1 cleaves 2’3’-c-di-AMP and 3’3’-cGAMP with comparable observed rates, while 2H PDE-9 is notably deficient at 2’3’-c-di-AMP degradation (Fig. 3A, B). This striking difference in the recognition of two highly similar substrates suggests that 2H PDE-9 may have adapted to evolutionary pressure to avoid 2’3’-c-diAMP.

Given this difference in activity, we investigated the structural relationship between Acb1 and 2H PDE-9. ColabFold predictions of 2H PDE-9 and Acb1 place them both as part of structural group 3, consistent with their similar enzymatic activity and broad-spectrum degradation of CBASS-related messengers. While structural prediction generates high confidence models of the enzyme core, the lid region is an important determinant of activity ^8^ and is not predicted with high reliability (Supplementary Fig. 9). We solved a 2.03 Å resolution structure of 2H PDE-9 (H50A) in the apo form and a 2.00 Å resolution 2H PDE-9 wild-type structure bound to a negatively charged ligand (citrate) (Fig. 3C, Supplementary Table 3, 4, Supplementary Fig. 10).

Previous structures of phage (PDB ID: 7T27) and eukaryotic viral (PDB ID: 9BKQ) PDEs have revealed that the 2H core employs a flexible lid that closes over the top of the captured ligand to enable phosphodiesterase activity ^5^. Phage Acb1 proteins use a short C-terminal lid domain which becomes ordered during nucleotide recognition. However, while the eukaryotic penguinpox virus PDE similarly shares conserved use of a lid, it spans from the opposite protein terminus ^8^. This variation was thought to be an example of convergent evolution that contributes to kingdom-specific adaptation toward nucleotide immune signals ^8^. Unexpectedly, although 2H PDE-9 and Acb1 exhibit moderate sequence similarity (44.8%) and fall into the same predicted structural group, the functional loop regions occur on different termini (Fig. 3D, E). The PDE-9 protein structure reveals an N-terminal lid that cascades over the active site in the presence of a negatively charged ligand, and adopts an ordered conformation along the outside of the protein in the apo form (Fig. 3D, Supplementary Fig. 11). This positional and structural divergence, coupled with conserved function, may reflect convergent evolution within phage or domain rearrangement and functional retention.

We next used TLC to determine if the lid is responsible for the substrate specificity of PDE-9. A deletion mutant lacking the lid (Fig. 3F) confirmed that the lid drives PDE-9 hydrolysis of 2’3’-c-di-AMP and 3’3’-cGAMP (Fig. 3G, Supplementary Fig. 12C). In a reaction with 3’3’-c-GAMP, the lid deletion was well tolerated by PDE-9, enabling efficient degradation over 1 hour even when the enzyme was lacking the 15 N-terminal residues (Fig. 3H, Supplementary Fig. 12A). In these conditions, the lid has a larger effect on enabling reaction with the less preferred substrate – and may be an adaptive tool toward increasing binding affinity for a broader range of oligonucleotides. Appending the T4 Acb1 lid to the C-terminus of PDE-9 lacking its N-terminal lid (Fig. 3F) fully restored activity, yet retained the substrate preference of PDE-9 (Fig. 3G, H, Supplementary Fig. 12A,B). Therefore, the preferential recognition of 3’3’-cGAMP by PDE-9 is not governed by the lid residues. These findings demonstrate the modularity of the PDE core and lid and indicate that substrate recognition is governed predominantly by the protein core rather than the structurally variable lid region. These results align with our structural classification, which reliably predicts groups of similar substrate specificity although lid domains are often modeled with low confidence in predictive structures (Supplementary Fig. 9).

### Viral 2H PDEs evade distinct immune systems

Immune pathways comprise nucleotide signals with varying chemical features. Signal diversity exists even within immune system types – notably, CBASS produces an array of signaling molecules including cGAMP, cUMP-AMP, and c-di-UMP ^9^. Chemical diversity enables host signaling fidelity and effector modularity and presents a challenge for viruses attempting to evade signaling. Acb1, the phosphodiesterase first found to protect from CBASS defense, is among the most promiscuous phosphodiesterases in our activity screen (Fig. 2D). Flexibility in substrate recognition allows Acb1 to combat different types of CBASS systems. However, this represents a potential liability if cell signaling or homeostasis is disrupted by viral PDE off-target activity.

Recently, we described a bacterial immune system, Panoptes, in which the role of the oligonucleotide messenger is reversed. In this system, 2’3’-c-di-AMP is synthesized independently of infection status and serves to repress – rather than activate – a toxic effector protein^20,21^. Panoptes preempts virus-mediated immune suppression, preventing the spread of viruses that contain enzymes such as Acb1 which deplete host cyclic nucleotides with limited specificity. Acb1 is an activator of the Panoptes due to its ability to recognize and degrade 2’3’-c-di-AMP.

Divergent substrate specificities may reflect adaptations to facilitate evasion of distinct immune pathways. Selective avoidance of 2’3’-c-di-AMP by PDE-9 suggested that it could evade Panoptes immunity. To test whether PDE-9 was sufficient for Panoptes activation, we coexpressed the Panoptes system and candidate phosphodiesterases (Fig. 4A). Strikingly, we found that induction of T4 Acb1 but not PDE-9 activated Panoptes-induced cell death (Fig. 4B, C). A deletion mutant of T4 Acb1 lacking the lid also did not activate cell death, confirming that the lid is necessary for catalysis (Fig 4D). However, a chimeric PDE-9 with a T4 Acb1 lid (Fig. 4D) did not activate cell death – this confirms our observation that the core of the protein is responsible for substrate specificity. The combined ability of PDE-9 to degrade CBASS messengers in combination with the selective avoidance of Panoptes repressor molecules demonstrates the ability of phage to evade multiple immune systems via a single antidefense protein.

## Discussion

2H PDEs represent a widespread family of virus-encoded enzymes capable of degrading diverse nucleotide messengers involved in cellular homeostasis and anti-viral immunity. These viral 2H PDEs are structurally diverse and, while they maintain the 2H core, this structural diversity is associated with the ability to cleave distinct sets of substrates. This has several important implications. First, rather than being restricted to acting as CBASS antagonists, we find that some 2H PDEs can degrade nucleotide messengers involved in other bacterial immune signalling pathways. Second, we find that this substrate specificity is associated with structural differences in the 2H core fold, raising the possibility that structural differences in the core 2H fold could be used to predict 2H PDE substrate affinity. Third, we find that differences in substrate specificity can yield 2H PDEs capable of degrading the CBASS signal 3’3’-cGAMP while avoiding the Panoptes signal 2’3’-c-diAMP, thus preventing cell death in the presence of Panoptes (Fig. 4E,F). Together, these data suggest that 2H PDEs are diverse and play a major role in the virus-host conflict.

We previously found that viruses that infect bacteria and viruses that infect animals encode 2H PDEs that can degrade cGAMP, which is produced by CD-NTases across the tree of life. Here, we expand that diversity of known viral 2H PDEs, and find that PDEs are capable of degrading diverse immune signals. We find that the 2H lid structure has likely arisen multiple times over the course of 2H PDE evolution. Despite high overall sequence identity, the Acb1 and 2H PDE-9 lids occur on opposite termini, suggesting that the 2H lid is a recent and recurrent adaptation in this lineage. Furthermore, we find that the lid of 2H PDE-9 is required for efficient hydrolysis of its preferred substrate, 3’3’-cGAMP, and is strictly required for activity toward its unpreferred substrate, 2’3’-cdiAMP. This suggests that the 2H lid may have a role in the adaptation of 2H PDEs against novel substrates.

The substrate preferences observed here may offer a roadmap to discovering new immune messengers or inhibitory molecules. Some of the studied PDEs do not cleave any of the nucleotides represented in this panel, raising the possibility of yet uncharacterized signals or immune systems. Novel signaling chemistries are still being uncovered – including the recent discovery that some Type III CRISPR systems use a unique SAM-AMP conjugate ^26^, base-modified (deoxyinosine) nucleotides as immune messengers ^27^, distinct ADPR analogs in TIR signaling ^28^, and the use of 2’3’c-diAMP as a repressive signaling molecule in Panoptes ^20,21^. Other PDEs exhibit broad substrate specificity with notable exceptions – for example, PDE-9 does not efficiently cleave 2’3’-c-di-AMP – suggesting that these omissions may reflect a selective pressure to preserve certain signaling molecules. Tailoring the PDE core to recognize only specific chemical messengers still comes at the cost of full spectrum evasion. Notably, other CBASS messengers, 2’3’-cGAMP and 3’3’cUAMP ^9,29^, are not efficiently degraded by PDE-9, which could be a casualty of evolution toward 2’3’-c-diAMP evasion.

Together, these findings suggest that variation in PDE substrate specificity reflects a dynamic evolutionary arms race, with both viruses and their hosts continuously redefining the signaling landscape. Recurrent adaptation of the 2H PDE core likely enables viruses to counter molecules responsible for a variety of bacterial immune defenses. Investigating these differences in PDE substrate preferences may reveal not only new signaling chemistries but also deepen our understanding of how specific molecular recognition shapes immune conflict.

## Acknowledgements

We thank members of the Doudna lab including Owen Tuck, Peter Yoon and Kenneth Loi for helpful discussion and feedback. We thank Keana Lucas for her exceptional operational and administrative leadership in support of the lab’s research and activities. J.A.D. is an investigator of the Howard Hughes Medical Institute, and research in the Doudna laboratory is supported by the Howard Hughes Medical Institute (HHMI). E.E.D. was supported by NIGMS of the NIH under award number F32GM153031. B.A.A. was supported by m-CAFEs Microbial Community Analysis and Functional Evaluation in Soils, a Science Focus Area led by Lawrence Berkeley National Laboratory based on work supported by the US Department of Energy, Office of Science, Office of Biological and Environmental Research under contract number DE-AC02-05CH11231. S.L. acknowledges support from the TED Audacious Fund. N.P. acknowledges support from the CRISPR Cures for Cancer Initiative. N.B. was supported by a University of California Office of the President funded UC-Historically Black Colleges and Universities Initiative (UC-HBC) award to the Doudna lab. We acknowledge the generous support of the James B. Pendleton Charitable Trust. Portions of this research were conducted on the Wynton Cluster at UCSF, supported by UCSF IT. Use of the Stanford Synchrotron Radiation Lightsource (SSRL), SLAC National Accelerator Laboratory, is supported by the U.S. Department of Energy, Office of Science, Office of Basic Energy Sciences under Contract No. DE-AC02-76SF00515. The SSRL Structural Molecular Biology Program is supported by the DOE Office of Biological and Environmental Research, and by the National Institutes of Health, National Institute of General Medical Sciences (P30GM133894). We thank Dr. Clyde Smith for expert assistance on beamline 12-2 at SSRL. This research used resources of the Advanced Light Source (ALS), a U.S. DOE Office of Science User Facility under contract no. DE-AC02-05CH11231. We thank Dr. Jay Nix for expert assistance at the Advanced Light Source beamline 8.2.1. We thank Dr. Anthony T. Iavarone for expert assistance in collecting mass spectrometry data. The QB3/Chemistry Mass Spectrometry Facility received National Institutes of Health support (grant number 1S10OD020062-01). The contents of this publication are solely the responsibility of the authors and do not necessarily represent the official views of NIGMS or NIH.

## Author contributions

E.E.D., J.N., and B.A.A. conceived of the project. E.E.D., J.N., B.A.A., S.L., K.H., N.P., and N.B. performed experiments. J.N. performed the bioinformatics experiments and analyses. E.E.D. performed biochemical experiments and analyses. E.E.D. and N.B. performed structural experiments. B.A.A., S.L., and K.H. performed bacterial assays. E.E.D., J.N., and J.A.D. wrote the original draft of the manuscript. All authors edited the manuscript and support its conclusions.

## Data availability

All code specific to this work can be found on Github here: 34. The methods refer to vpSAT, which can be found here: https://github.com/jnoms/vpSAT. The structures and sequences of the 2H PDEs studied here, as well as a copy of the github repository 2025_Doherty-Nomburg-2H_PDEs, can be found on Zenodo with the following DOI: 10.5281/zenodo.16709871. The structure of PDE-9 H50A and PDE-9 wild-type have been deposited under PDB ID: 9Q2Y and PDB ID: 9Q2G respectively and will be released upon publication of the primary citation.

## Materials and Methods

### Structure prediction, alignments, and phylogenetics of 2H PDEs

MSAs were generated for phage proteins using local colabfold (colabfold_search) against the colabfold database, downloaded May 31 2023. Structure prediction used Colabfold ^12^ version 1.5.2, with the following settings: –num_recycle 3, –num_models 3, –use-gpu-relax, – stop_at_score 80, –stop_at_score_below 40. The model with the highest pLDDT was selected for use.

Structural alignments used DaliLite ^15^ version 5 against the database of predicted structures from phage, using predicted structures of the following known viral 2H PDEs as queries: T4 Acb1 (NP_049750), Iridovirus PDE (YP_009021100), MERS NS4b (YP_009047207), Pigeonpox PDE (YP_009046269), Porcine Torovirus ORF1ab (YP_008798230, 2H region only), Marsiellevirus PDE (YP_003406995), Rotavirus A VP3 (YP_002302228, 2H region only), and Murine hepatitis virus NS2a (YP_009824980). Alignments used vpSAT’s dali.sh. Proteins that aligned to a query with a DALI Z score of 5 or higher were manually assessed for conservation of two catalytic histidines and the overall 2H architecture, resulting in a final count of 75 2H PDEs.

Jackhmmer searches used HMMER version 3.1b2, with three recycles, to search the same query viral 2H PDEs against all phage protein sequences. Alignments with an evalue less than 0.001 were kept.

For phylogenetic analysis, a MSA was generated using Foldmason ^31^ version 1.763a428, using predicted structures of the known and novel viral 2H PDEs as inputs. For the 2H PDEs with multiple domains beyond the 2H, only the 2H domain was input into Foldmason. The MSA was used as input for iqtree version 1.6.12, using automatic model selection with the following settings: -m TEST -B 1000 --seqtype AA -T AUTO. The +I+G4 model was used for the resultant tree.

Domains associated with each phage 2H PDE were identified in the following manner. First, chainsaw ^32^ was used to split each 2H PDE into constituent domains, and 2H PDE domains were identified as domains that aligned with T4 Acb1 using DALI. The other, non-2H domains were aligned with the S20 nonredundant domains from CATH ^7^ (cath-dataset-nonredundant-S20.pdb.tgz), downloaded August 16 2024, using foldseek easy-search (version 427df8a6b5d0ef78bee0f98cd3e6faaca18f172d) with the setting -tmscore-threshold 0.5, and alignments with an alignment length above 60 residues were kept.

### Protein expression and purification

Proteins were expressed as purified previously described with the following modifications ^7^. In brief: Expression sequences for candidate phosphodiesterases were cloned into a custom pET-based vector by Gibson assembly to yield an N-terminal His_10_-MBP-TEV construct. For single domain proteins, the entire coding sequence was used in expression. For multi-domain proteins (2H PDE-13, 2H PDE-16) only the 2H domain was cloned into the plasmid for expression.

Proteins were expressed in *E. coli* Rosetta 2 (DE3) pLysS by growing cells to an OD_600_ of 0.4-0.6 in 2xYT (2x yeast extract tryptone) medium at 37°C, and induced with 0.5 mM IPTG (Isopropyl β-D-1-thiogalactopyranoside) following a cold shock at 4°C. After induction, cells expressing each protein were grown overnight at 16°C. Cells were collected by centrifugation for 20 min at 4,000 rpm, 4°C and resuspended in 20 mM Tris-HCl, pH 8.0, 10 mM imidazole, 2 mM MgCl_2_, 500 mM KCl, 10% (v/v) glycerol, 0.5 mM Tris (2-carboxyethyl) phosphine, and Roche protease inhibitor.

Cells were lysed by sonication, and cell lysate was clarified by centrifugation at 17,000 x g, 4 °C for 0.5 h. The supernatant was bound to Nickel-NTA affinity resin pre-equilibrated with wash buffer (20 mM Tris-HCl, pH 8.0, 500 mM KCl, 30 mM imidazole, 10% (v/v) glycerol, and 0.5 mM Tris(2-carboxyethyl) phosphine) for 1 h at 4°C. Supernatant was discarded and resin was washed 5 × 30 mL wash buffer (20 mM Tris-HCl, pH 8.0, 500 mM KCl, 30 mM imidazole, 10% (v/v) glycerol, and 0.5 mM Tris(2-carboxyethyl) phosphine). Protein was eluted in 5 mL elution buffer (20 mM Tris-HCl, pH 8.0, 500 mM KCl, 300 mM imidazole, 10% (v/v) glycerol, and 0.5 mM Tris(2carboxyethyl) phosphine). Proteins for *in vitro* thin layer chromatography analysis were concentrated to 200 µM, buffer exchanged into 20 mM TrisHCl, pH 8.0, 500 mM KCl, 30 mM imidazole, 10% (v/v) glycerol, and 0.5 mM Tris(2-carboxyethyl) phosphine), snap-frozen and stored at −80°C.

For structural studies of 2H PDE-9, after nickel elution recombinant TEV Protease with an N-terminal His-tag was added to the elution for cleavage and dialyzed overnight at 4 °C in a 4,000 MWCO dialysis cassette (Thermo Fisher Scientific) with dialysis buffer (20 mM Tris-HCl, pH 8.0, 500 mM KCl, 30 mM imidazole, 10% (v/v) glycerol, and 0.5 mM Tris(2-carboxyethyl) phosphine). The resultant solution was passed over a 5 mL Ni-NTA Superflow cartridge (Cytiva). Flow-through was collected, concentrated using a MWCO centrifugal filter (Amicon), and loaded onto a HiLoad 16/600 Superdex 200 pg column (Cytiva). Elution was isocratic (20 mM Tris-HCl, pH 8.0, 500 mM KCl, 10%, 1 mM TCEP) glycerol monitored by A280, peaks were pooled, concentrated to 15 mg/mL, snap-frozen, and stored at −80°C.

### Analysis of recombinant protein

Purified protein was analyzed by SDS-PAGE. Samples were prepared in 1X Protein Loading Dye (50 mM Tris-HCl, pH 6.8, 15 mM EDTA, 6% (v/v) glycerol, 10% SDS, and bromophenol blue), heated at 95 °C for 3 min, loaded onto a 12% Mini-Protean TGX Precast Protein Gel (Bio-Rad) and run at 125 V until the dye front reached the bottom of the gel. Gels were stained in 30% ethanol, 10% glacial acetic acid in water, 0.1% (w/v) R-250 Coomassie and destained in 40% ethanol, 10% glacial acetic acid in water.

### Phosphodiesterase activity reactions

Recombinant enzymes were assayed for phosphodiesterase activity by in vitro reactions with an array of oligonucleotides. All compounds were purchased from Biolog Life Science Institute and Enzo Life Sciences and used without further purification. Reactions containing recombinant phosphodiesterases and candidate molecules were assayed as previously described, with the following modifications ^33^ .Briefly, reactions were initiated by the addition of 40 µM recombinant enzyme in reaction buffer (50 mM Tris, pH 8.0, 10 mM MgCl_2_,100 mM NaCl) to 1.25 mM oligonucleotide. Each substrate was also run with the exclusion of enzyme and with the inclusion of an unrelated, non-phosphodiesterase enzyme (AFY13309.1) purified in the same conditions. The reaction mixture was incubated at 37°C for 5 minutes and stopped by heating to 95°C for 2 min. Reactions were performed in technical duplicate.

### Thin layer chromatography (TLC)

Silica gel TLC plates L × W 5 cm × 10 cm with fluorescent indicator 254 nm were spotted with 2 µL in vitro enzymatic reaction. Separation was performed in an eluent of n-propanol/ammonium hydroxide/ water (11:7:2 v/v/v). The plate was allowed to dry fully and visualized with a short-wave ultraviolet light source at 254 nm.

### Quantification of Product formation by TLC

Reactions were initiated as described above at 8 µM recombinant enzyme in reaction buffer (50 mM Tris, pH 8.0, 10 mM MgCl_2_,100 mM NaCl) with 1.25 mM oligonucleotide. The reaction mixture was incubated at 37°C for the time indicated and stopped by heating to 95°C for 2 min. Thin layer chromatography was performed as indicated above and visualized with a short-wave ultraviolet light source at 254 nm. Non-degraded substrate spots were defined by comparison to a chemical standard. The ratio of substrate to degradation products for each timepoint was determined using densitometry of TLC images in Gel Analyzer 19.0 using a rolling ball background detection parameter with a 30% peak width tolerance. Percent degradation was determined by dividing the intensity of the substrate band by the intensity of the background for each lane. Values were normalized with the negative control set to 0% degradation. For kinetic measurements, data were fit to a one phase decay equation (Y=Y0 - Plateau)*exp(−K*X) + Plateau) in Graphpad Prism 10.1.0. Values plotted are the mean of three technical replicates and error bars represent the standard deviation.

### Crystallization and structure determination

Crystals of 2H PDE-9 wild-type or H50A (9Q2G, 9Q2Y) in the closed lid or apo/open lid form respectively were grown at 20 °C using the hanging drop vapor diffusion method.

For 2H PDE-9 H50A crystals in the apo form, a 1 µL solution of 3 mg/mL 2H PDE-9 H50A in 20 mM Tris-HCl, pH 8.0, 500 mM NaCl, 5% glycerol, and 2.5 mM 3’3’-cGAMP was mixed with 1 µL 0.2 M magnesium acetate tetrahydrate, 0.1 M sodium cacodylate trihydrate, pH 6.5, and 20% w/v polyethylene glycol (PEG) 8,000. Single crystals appeared within 3 days and were cryoprotected in a solution of mother liquor with 30% glycerol before being flash cooled in liquid nitrogen.

For 2H PDE-9 wild type crystals in the closed lid form (citrate bound in active site), 1 µL solution of 3 mg/mL 2H PDE-9 in 20 mM Tris-HCl, pH 8.0, 500 mM NaCl, 5% glycerol, and 2.5 mM 3’3’-cGAMP was mixed with 1 µL 0.2 M ammonium acetate, 0.1 M sodium citrate tribasic dihydrate, pH 5.6, 30% PEG 4,000 and supplemented with 0.2 µL 10 mM phosphorothioate modified 3’3’-cGAMP (c[A(3’,5’)pS-G(3’,5’)pS “isomer 2” from Biolog). Single crystals appeared within 1 day and were cryoprotected in a solution of mother liquor with 30% glycerol before being flash cooled in liquid nitrogen.

Data for 2H PDE-9 H50A in the apo/open lid form were collected *via* fine-phi slicing using 0.2° oscillations at beamline 8.2.1 at the Advanced Light Source at Lawrence Berkeley National Laboratory. X-ray diffraction data were measured to 2.03 Å.

Data for 2H PDE-9 wild-type in the closed lid form were collected *via* fine-phi slicing using 0.2° oscillations at beamline 12-2 at Stanford Synchrotron Radiation Lightsource at SLAC National Accelerator Laboratory. X-ray diffraction data were measured to 1.74 Å.

### Processing and refinement of crystallographic data

Crystallographic data were processed in CCP4i2 ^34^ with xia2 ^35^ using DIALS for indexing, refinement, and integration using POINTLESS ^36^ and AIMLESS ^30^ for scaling. For 2H PDE-9 H50A, a resolution cutoff of 2.00 Å was applied to scaling in AIMLESS to generate a robust complete dataset. The structures were solved by molecular replacement using a Colab-Fold ^37^ generated model of 2H PDE-9 wild-type or mutated to H50A in Sculptor (Phenix) ^37^.

Molecular replacement successfully identified the placement of one wildtype PDE-9 molecule as indicated by the log-likelihood gain (LLG) 5729 and the translation-function Z-score (TFZ) 64.5. Molecular replacement successfully identified the placement of two PDE-9 H50A monomers as indicated by the log-likelihood gain (LLG) 925 and the translation-function Z-score (TFZ) 32.4. The structures were refined using PHENIX ^21^ including simulated annealing, non-crystallographic symmetry, and TLS parameters. Each structure contained disordered regions where residues could not be modeled into the density (PDE-9 wild type residue 1, PDE-9 H50A residues 1-3, 165). Both models were built and adjusted using COOT. The H50A PDE-9 structure was refined to a final R_free_ and R_work_ of 24.49% and 22.58% respectively (**Supplementary Table 3**). The wild-type PDE-9 structure was refined to a final R_free_ and R_work_ of 19.6% and 14.77% respectively (**Supplementary Table 4**). Atomic coordinates and structures factors have been deposited to the Protein Data Bank.

### Mass spectrometry sample preparation

Reactions between phosphodiesterase and oligonucleotides as described above were diluted 1:2 in deionized H_2_O, centrifuged at 13,000 x g for 15 min and used directly in subsequent analysis.

### Mass spectrometry

Oligonucleotide degradation products were analyzed using a liquid chromatography (LC) system (1200 series, Agilent Technologies, Santa Clara, CA) as previously described ^38^. The LC system was connected in line with an LTQ-Orbitrap-XL mass spectrometer equipped with an electrospray ionization (ESI) source (Thermo Fisher Scientific, Waltham, MA). The LC system was equipped with a G1322A solvent degasser, G1311A quaternary pump, G1316A thermostatted column compartment, and G1329A autosampler unit (Agilent). The column compartment was equipped with an Ultra C18 column (length: 150 mm, inner diameter: 2.1 mm, particle size: 3 µm, catalog number: 9174362, Restek, Bellefonte, PA). Ammonium acetate (≥98%, Sigma-Aldrich, St. Louis, MO), methanol (Optima LC-MS grade, 99.9% minimum, Fisher, Pittsburgh, PA) and water purified to a resistivity of 18.2 MΩ·cm (at 25 °C) using a Milli-Q Gradient ultrapure water purification system (Millipore, Billerica, MA) were used to prepare mobile phase solvents. Mobile phase solvent A was water and mobile phase solvent B was methanol, both of which contained 10 mM ammonium acetate. The elution program consisted of isocratic flow at 0.5% (volume/volume) B for 2 min, a linear gradient to 99.5% B over 2 min, isocratic flow at 99.5% B for 4 min, a linear gradient to 0.5% B over 1 min, and isocratic flow at 0.5% B for 21 min, at a flow rate of 100 µL/min. The column compartment was maintained at 30 °C and the sample injection volume was 1 µL. External mass calibration was performed in the positive ion mode using the Pierce LTQ ESI positive ion calibration solution (catalog number 88322, Thermo Fisher Scientific). Full-scan, high-resolution mass spectra were acquired in the positive ion mode over the range of mass-to-charge ratio (*m*/*z*) = 300 to 2000, using the Orbitrap mass analyzer, in profile format, with a mass resolution setting of 60,000 (at *m*/*z* = 400, measured at full width at half-maximum peak height, FWHM). Tandem mass (MS/MS or MS^2^) spectra were acquired using collision-induced dissociation (CID) in the linear ion trap, in centroid format, with the following parameters: isolation width = 3 *m*/*z* units, normalized collision energy = 28%, activation Q = 0.25, and activation time = 30 ms. Data acquisition was controlled using Xcalibur software (version 2.0.7, Thermo Fisher Scientific).

### Mass spectrometry data processing

Raw data were converted to mzXML format using msconvert 3.0.19052.1 from the Galaxy platform ^21^. Data were then processed using the open source software MZmine 3.9.0. Compound identification was performed by comparing retention times and mass-to-charge ratios (*m*/*z*) with those of chemical standards.

### Panoptes activator assays

To test whether Panoptes was activated by different viral 2H PDEs, we cloned Panoptes in its native context into a p15a-CmR plasmid and candidate activators under control of the pJex promoter in a low copy SC101-KanR plasmid. dCas13d under pTet on a p15a-CmR plasmid (pBA635) as employed as Panoptes negative control ^21^. RFP under pJex on a SC101-KanR plasmid was employed as an activator negative control.

To perform liquid culture activator assays, 3 independent overnight cultures containing candidate Panoptes and candidate activator plasmids were inoculated in LB media at 8e6 CFU in a Corning 3903 plate with the following supplements: 35 µg/mL Chloramphenicol, 50 µg/mL Kanamycin, variable amounts of crystal violet (CV) to induce candidate activator expression and no aTc for Panoptes expression. For candidate activator expression conditions, +250 nM CV was used. The plate was monitored in a Cytation5 plate reader (Biotek) at 30°C, 807 rpm and OD600 measured every 5 min for 12 h. Data were plotted using the Seaborn package in Python.

