## Supplementary Figures for "Divergent viral phosphodiesterases for immune signaling evasion"

**Supplementary Fig. 1. Target proteins detected by different alignment methods.**
**Supplementary Fig. 2. TLC method schematic.**
**Supplementary Fig. 3. Representative TLCs from in vitro degradation reactions.**
**Supplementary Fig. 4. Complete heatmap showing degradation of all PDEs vs. oligonucleotide panel.**
**Supplementary Fig. 5. Degradation product identification overview.**
**Supplementary Fig. 6. HPLC-MS identification of degradation products.**
**Supplementary Fig. 7. Detection of a shared intermediate by HPLC-MS.**
**Supplementary Fig. 8. Degradation of 2’3’c-diAMP and 3’3’-cGAMP by 2H PDE-9 and T4 Acb1 over time.**
**Supplementary Fig. 9. pLDDT of 2H PDE-9 prediction by AlphaFold 2.**
**Supplementary Fig. 10. Purification and crystallization of 2H PDE-9 wildtype and H50A.**
**Supplementary Fig. 11. Placement of negatively charged ligands in T4 Acb1 structure vs. PDE-9 structure.**
**Supplementary Fig. 12. Representative TLCs for quantification of substrate degradation in vitro.**

**Supplementary Tables**

**Supplementary Table 1. Manual curation of 2H PDE hits with intact catalytic histidines.**
**Supplementary Table 2. 2H PDEs with putative additional domains.**
**Supplementary Table 3. X-ray data collection and refinement statistics for PDB ID:** 9Q2G
**Supplementary Table 4. X-ray data collection and refinement statistics for PDB ID: 9Q2Y**

A

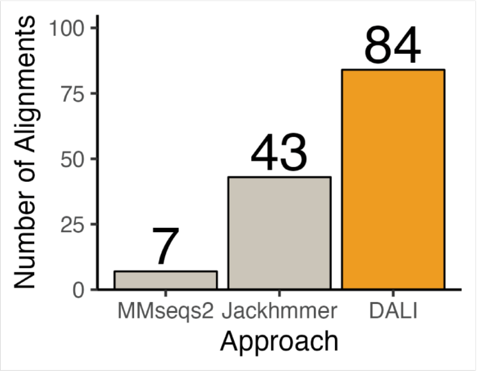


**Supplementary Fig. 1. Target proteins detected by different alignment methods. A)** We aligned the same panel of 8 known viral 2H PDEs against a set of phage proteins using MMseqs2 (a sequence alignment method), Jackhmmer (a HMM-based alignment method), and DALI (a structural alignment method). The number of target proteins detected by each method are indicated on the Y axis, with the approach indicated on the X axis.

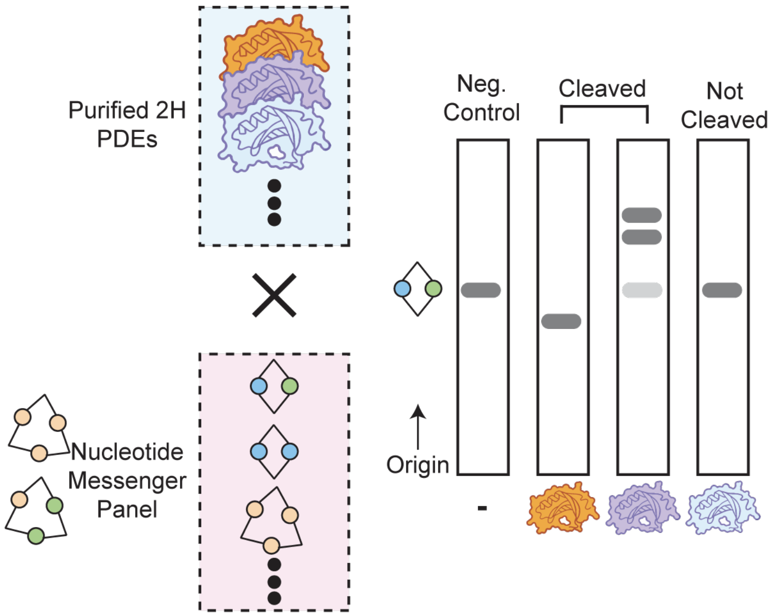


**Supplementary Fig. 2. TLC method schematic.** A panel of nucleotide messengers is subjected to reaction by different purified 2H PDEs. Each messenger is run by TLC and compared to the reaction of the nucleotide with each PDE. The presence of additional spots indicates cleavage, where partial cleavage still shows a spot that migrates consistently with the messenger molecule alone.

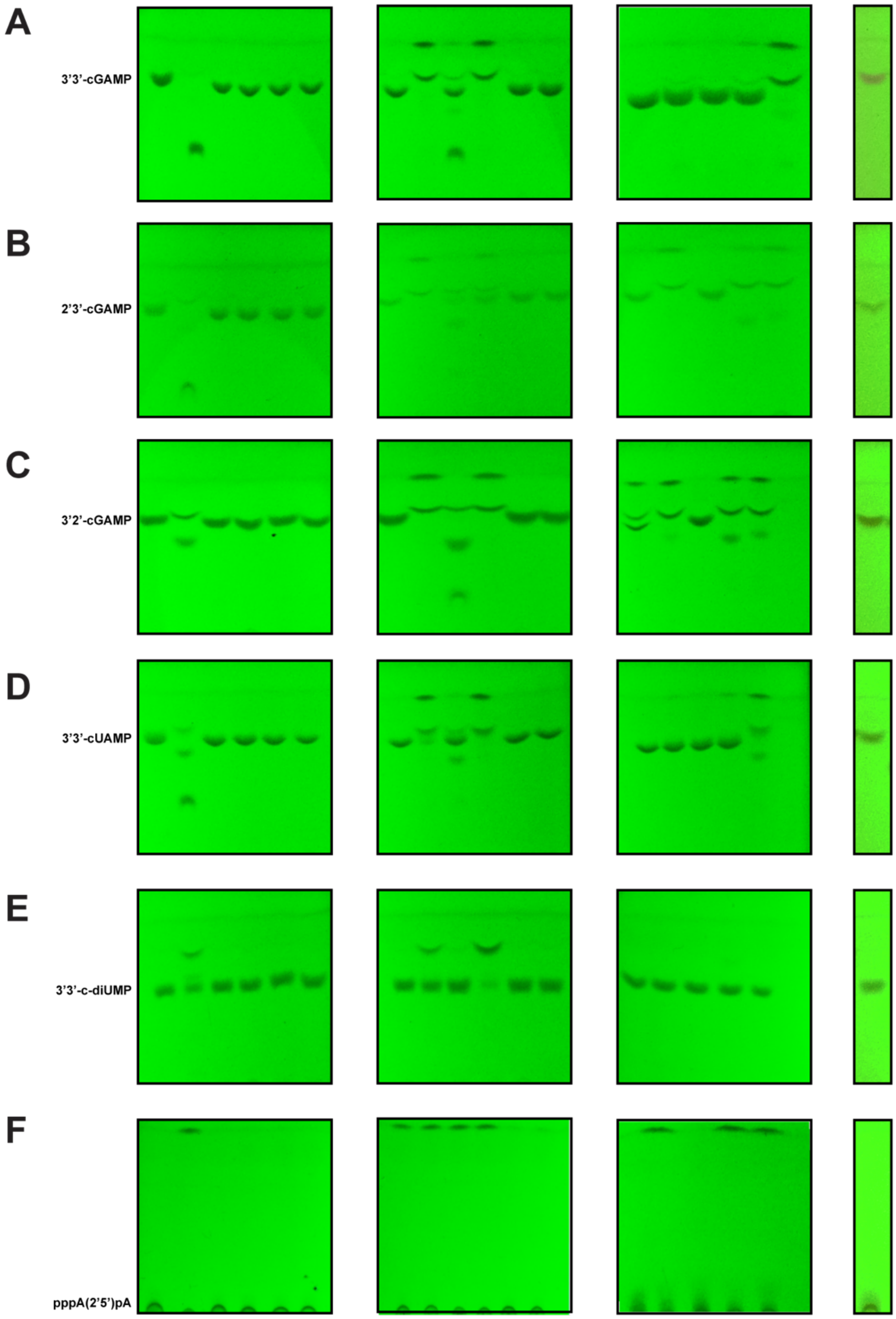
A
3'3'-cGAMP
B
2'3'-cGAMP
C
3'2'-cGAMP
D
3'3'-cUAMP
E
3'3'-c-diUMP
F
pppA(2'5')pA

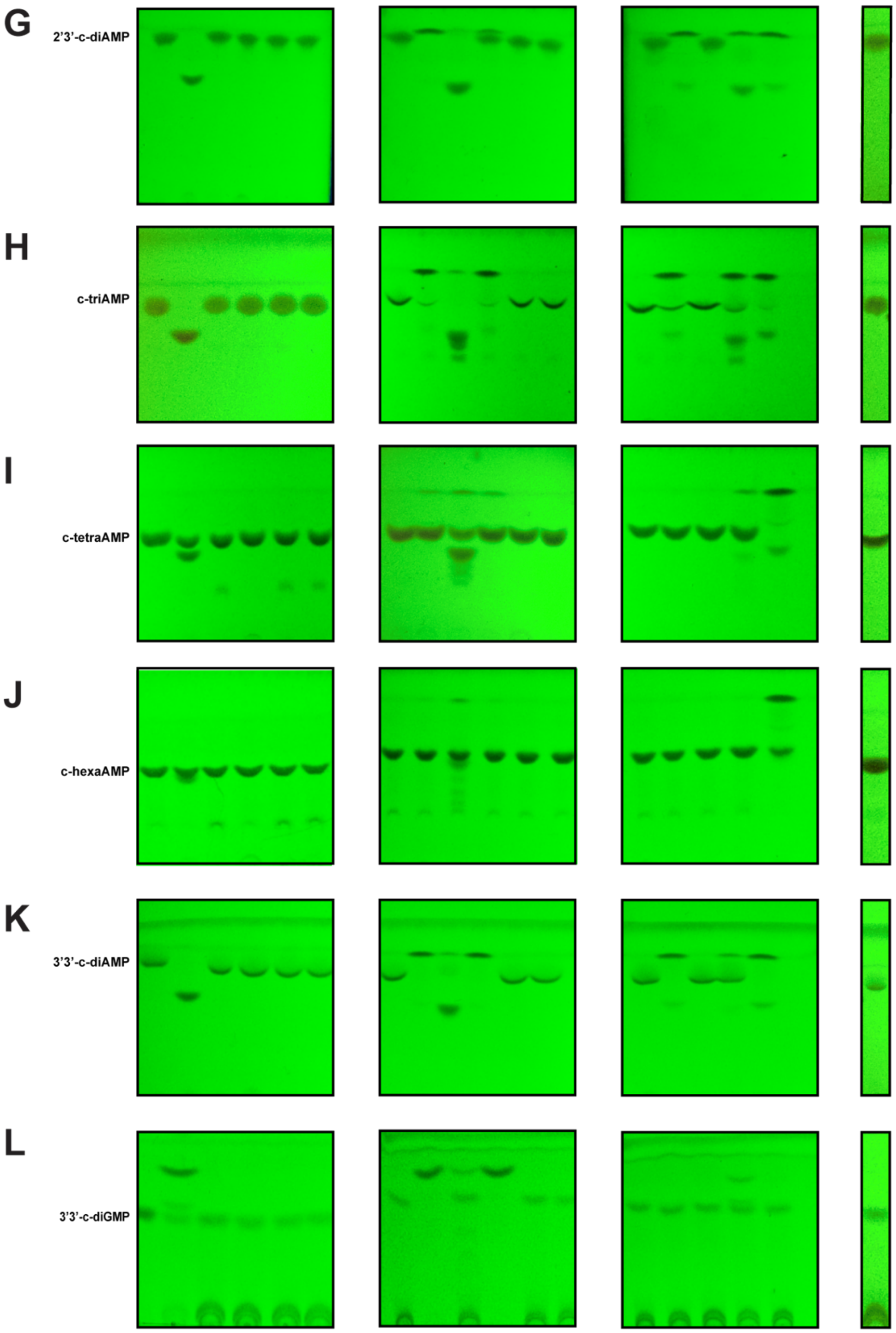
G
2'3'-c-diAMP
H
c-triAMP
I
c-tetraAMP
J
c-hexaAMP
K
3'3'-c-diAMP
L
3'3'-c-diGMP

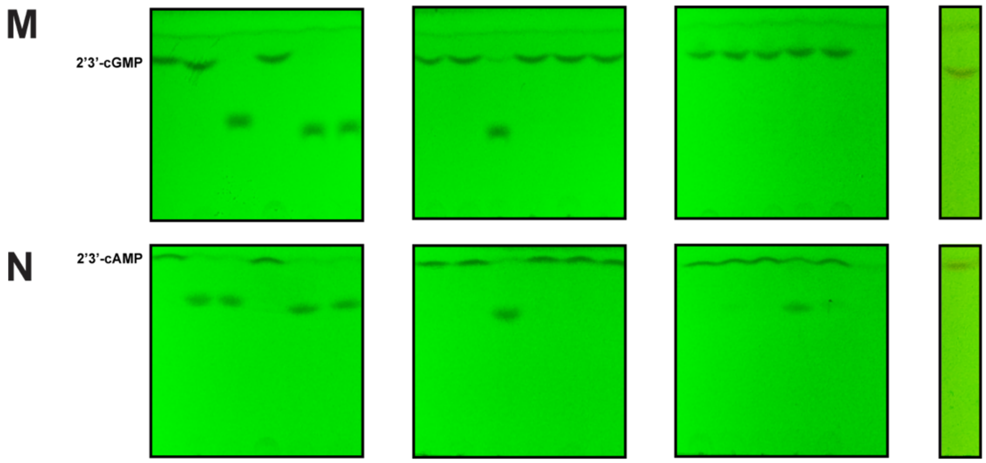


**Supplementary Fig. 3. Representative TLCs from in vitro degradation reactions.** Each reaction was completed and assessed by TLC in technical duplicate. Here are representative TLC plates for the reaction between a panel of PDEs (lanes in order are no enzyme, T4 Acb1, and then PDE-1, 3, 4, 5, 6, 7, 8, 9, 10 ,11, 12, 13, 14, 15, 16 and negative control enzyme (Methods)) and various nucleotides, where: **A)** 3'3'-cGAMP **B)** 2'3'-cGAMP **C)** 3'2'-cGAMP **D)** 3'3'-cUAMP **E)** 3'3'-c-diUMP **F)** pppA(2'5')pA **G)** 2'3'-c-diAMP **H)** c-triAMP **I)** c-tetraAMP **J)** c-hexaAMP **K)** 3'3'-c-diAMP **L)** 3'3'-c-diGMP **M)** 2'3'-cGMP **N)** 2'3'-cAMP.

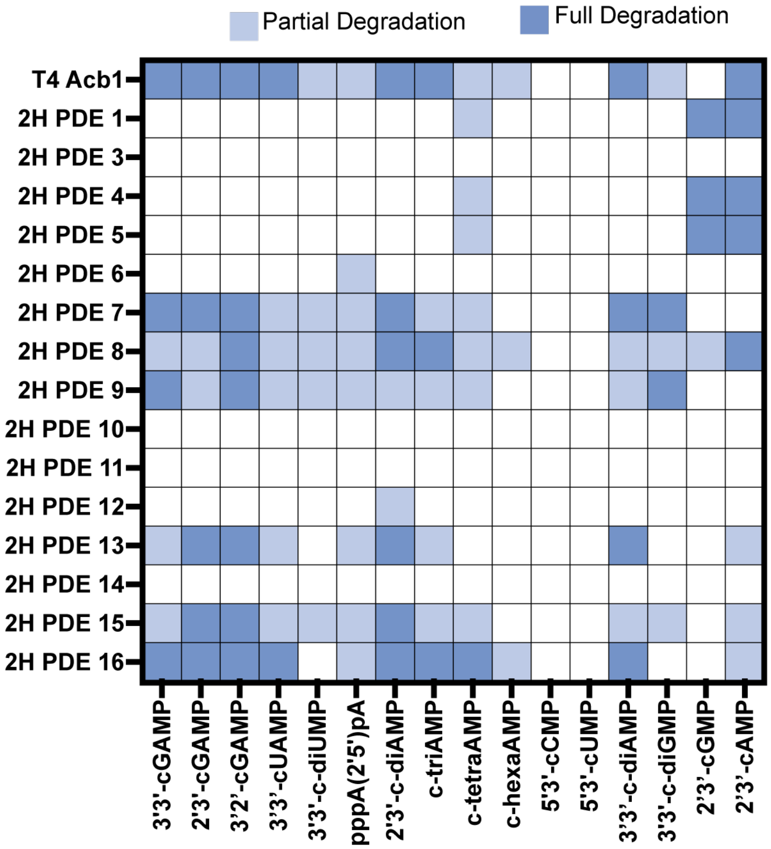


**Supplementary Fig. 4. Complete heatmap showing degradation of all PDEs vs. oligonucleotide panel.** Full or partial degradation indicated by light or dark blue for combinations of PDE and panel molecule.

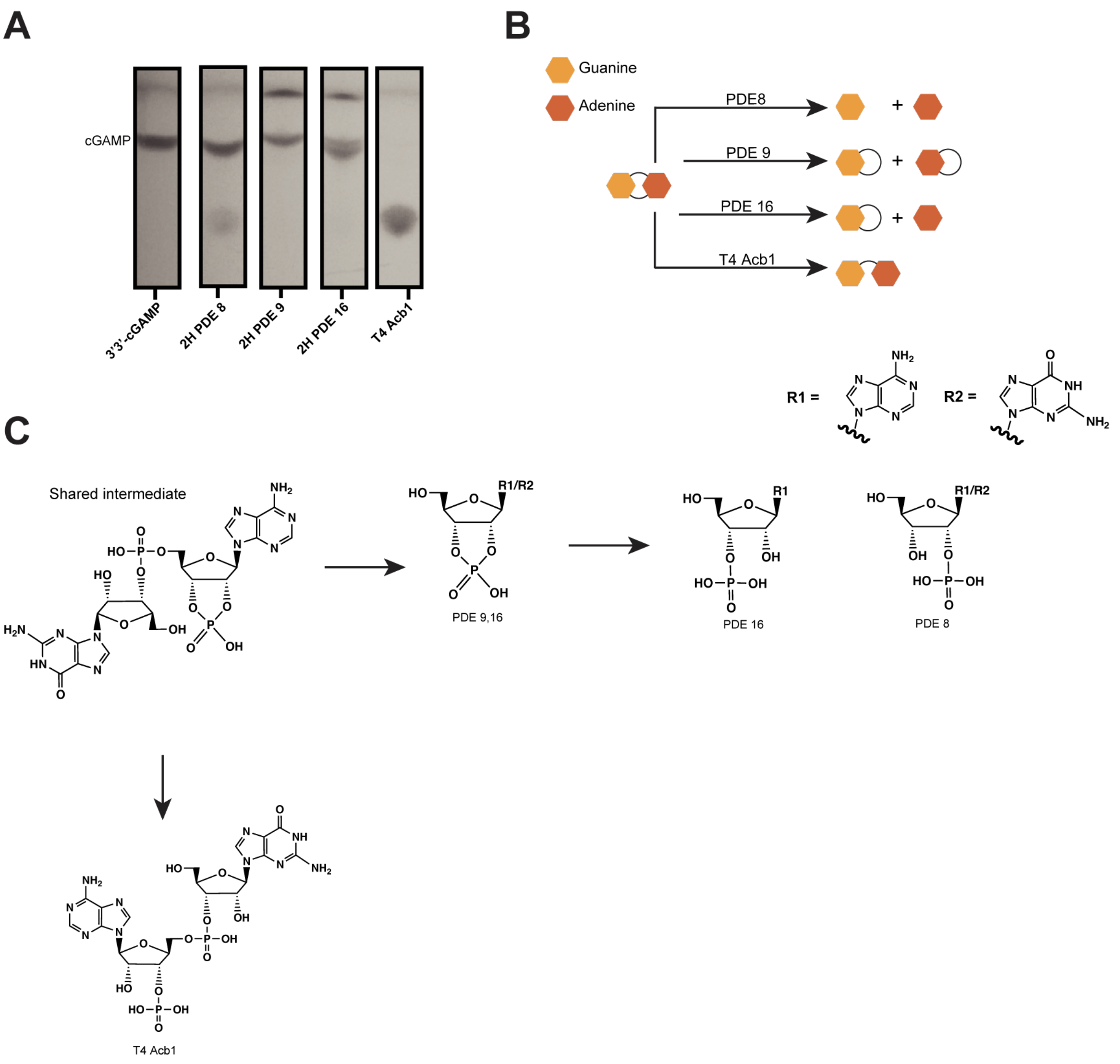


**Supplementary Fig. 5. Degradation product identification overview. A)** TLCs of in vitro reactions between 3'3'-cGAMP (labelled band) and various PDEs show the formation of different degradation products. **B)** Overview of products made by each PDE (Supplementary Fig. 6). **C)** Proposed mechanistic steps for the generation of different degradation products from a shared intermediate (Supplementary Fig 7).

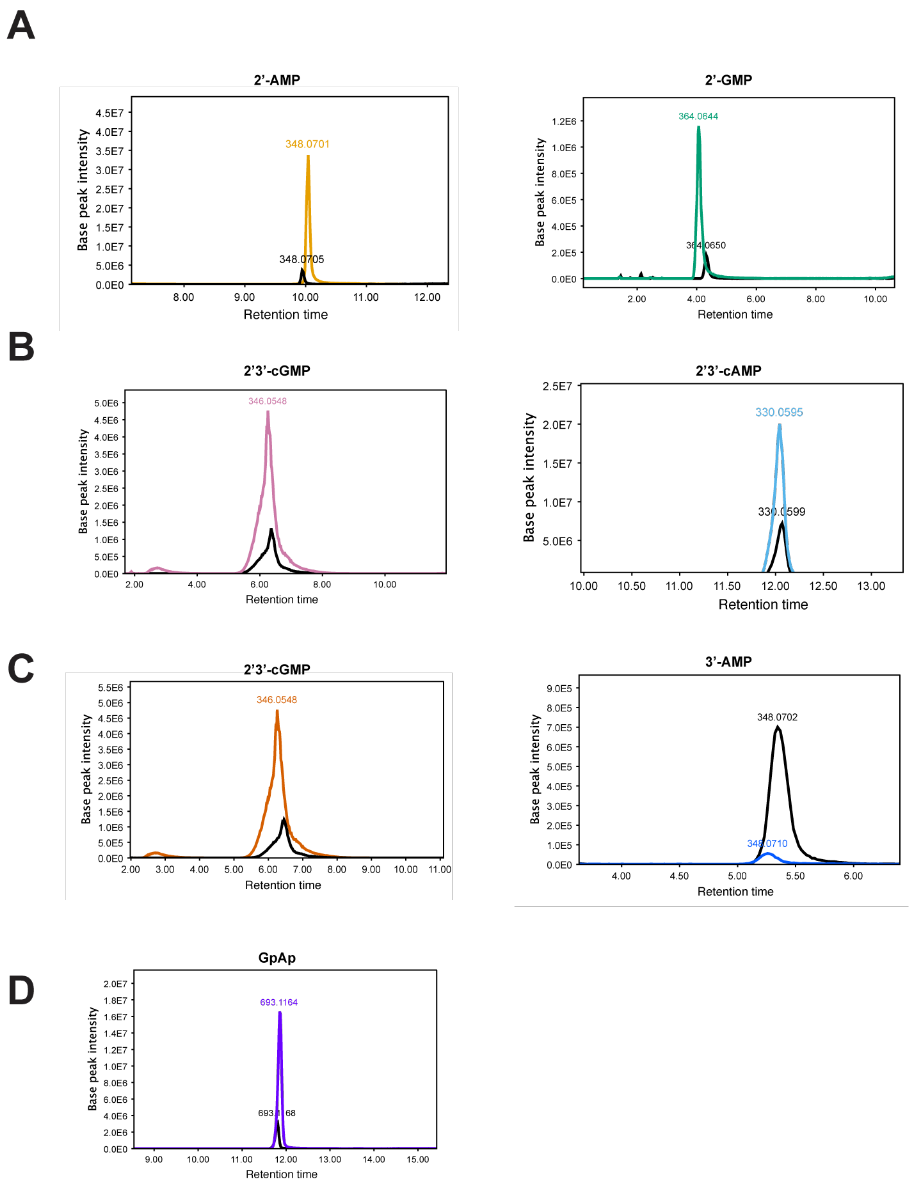


**Supplementary Fig. 6. HPLC-MS identification of degradation products.** Overlay of HPLC-MS traces of chemical standards versus PDE x 3'3'cGAMP reactions. **A)** Detection of products from reaction with 2H PDE-8. **B)** Detection of products from reaction with 2H PDE-9. **C)** Detection of products from reaction with 2H PDE-16. **D)** Detection of products from reaction with T4 Acb1.

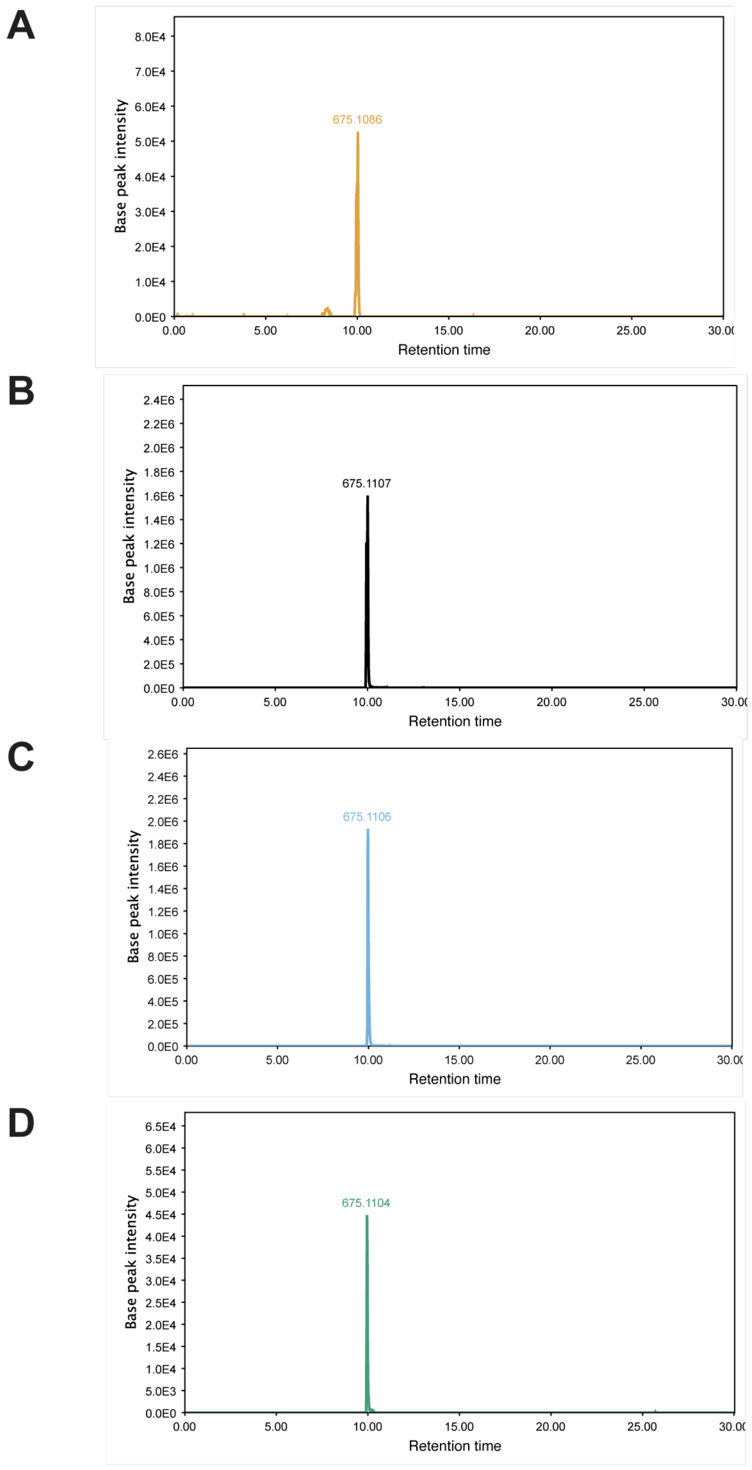


**Supplementary Fig. 7. Detection of a shared intermediate by HPLC-MS.** Extracted ion chromatograms (m/*z* = 675-676) for the reaction between 3'3'-cGAMP and 2H PDEs showing a molecular weight consistent with a GpA>p intermediate. **A)** 2H PDE-8. **B)** 2H PDE-9. **C)** 2H PDE-16. **D)** T4 Acb1.

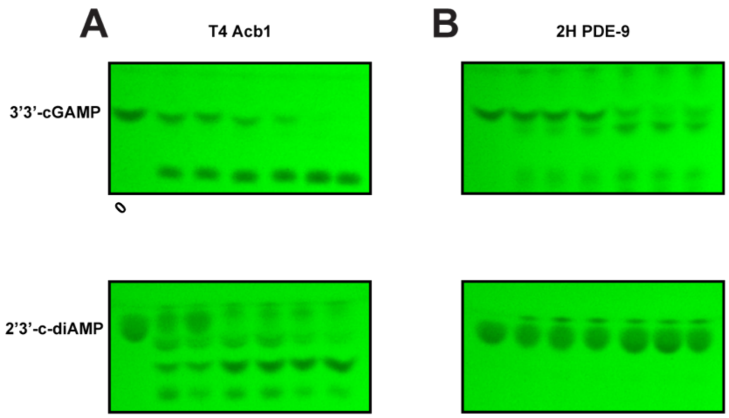


**Supplementary Fig. 8. Degradation of 2'3'c-diAMP and 3'3'-cGAMP by 2H PDE-9 and T4 Acb1 over time.** Representative TLCs used to quantify substrate over time as 3'3'-cGAMP and 2'3'-c-diAMP are consumed by reaction with **A)** T4 Acb1 and **B)** 2H PDE-9. Reactions were performed in technical triplicate.

A

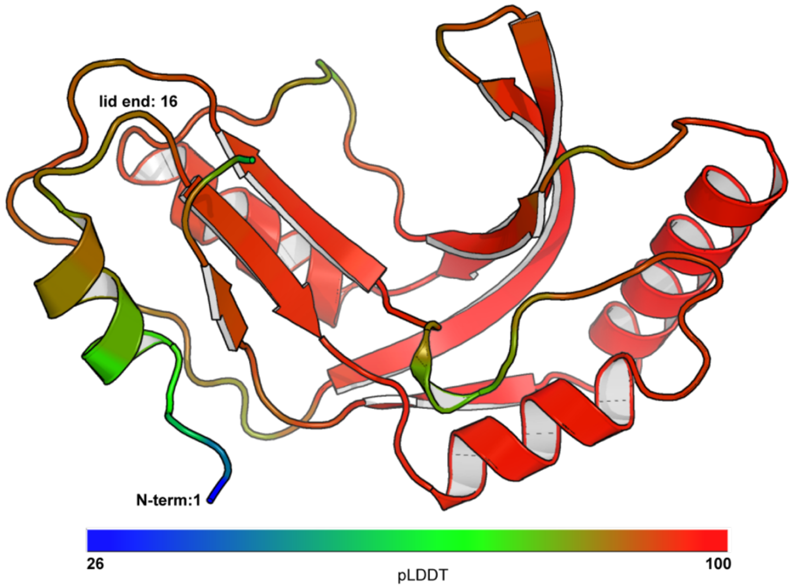


**Supplementary Fig. 9. pLDDT of 2H PDE-9 prediction by AlphaFold 2. A)** Predicted structure of 2H PDE colored by pLDDT (predicted local difference distance test; scale bar below structure) score where a pLDDT of 70 indicates correct backbone placement–sidechains may be misplaced–and 90 indicates a high confidence structure. The lid region is labeled spanning from the N-term (residue 1) to the end of the lid (residue 16).

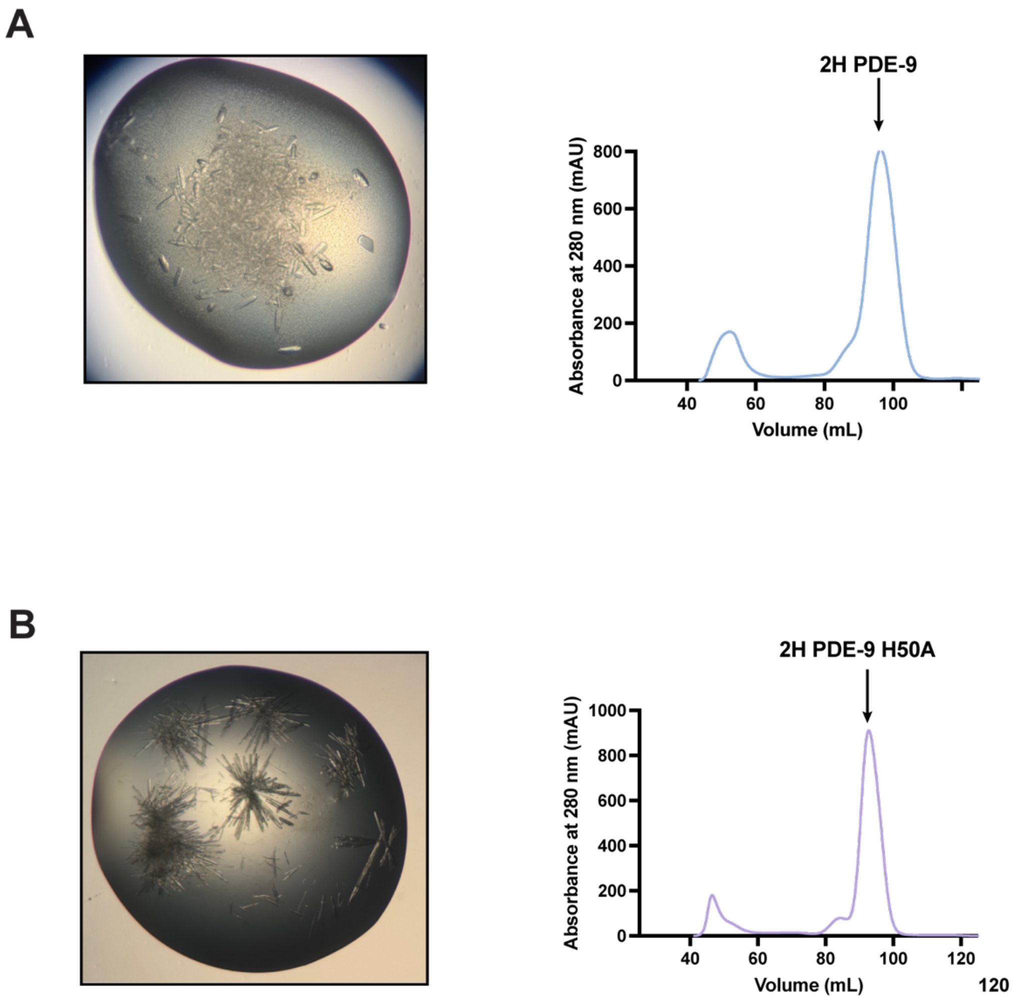


**Supplementary Fig. 10. Purification and crystallization of 2H PDE-9 wildtype and H50A. A) Left:** 2H PDE-9 wild-type crystals grown in 0.2 M ammonium acetate, 0.1 M sodium citrate tribasic dihydrate, pH 5.6, 30% PEG 4,000. **Right:** Size exclusion chromatography trace from purification of 2H PDE-9 indicates a monomeric species. The arrow indicates the peak collected for crystal trials. **B) Left:** 2H PDE-9 H50A crystals grown in 0.2 M magnesium acetate tetrahydrate, 0.1 M sodium cacodylate trihydrate, pH 6.5, and 20% w/v polyethylene glycol (PEG) 8,000. **Right:** Size exclusion chromatography trace from purification of 2H PDE-9 indicates a monomeric species. The arrow indicates the peak collected for crystal trials.

A

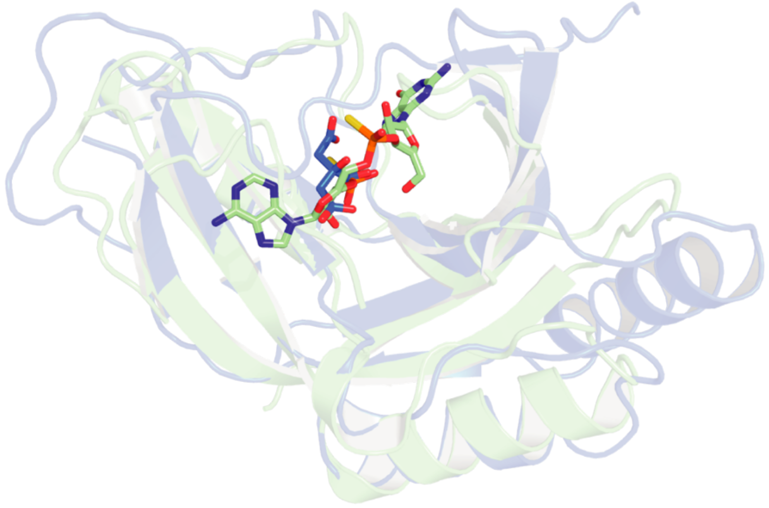

B

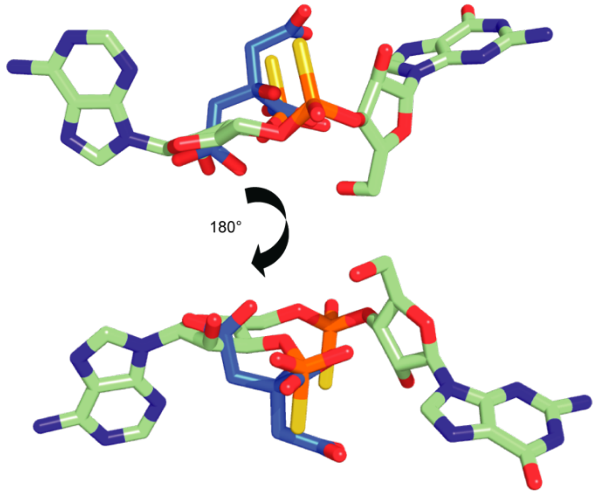


**Supplementary Fig. 11. Placement of negatively charged ligands in T4 Acb1 structure vs. PDE-9 structure. A)** Overlay of T4 Acb1 ligand bound structure (PDB ID: 7T27, green) with closed lid/ citrate bound structure of 2H PDE-9 (PDB ID: 9Q2G, blue). **B)** Overlay of citrate with the linearized 3’3’-cGAMP molecule (GpAp) shows a consistent positioning of the negatively charged citrate molecule and the phosphate of GpAp.

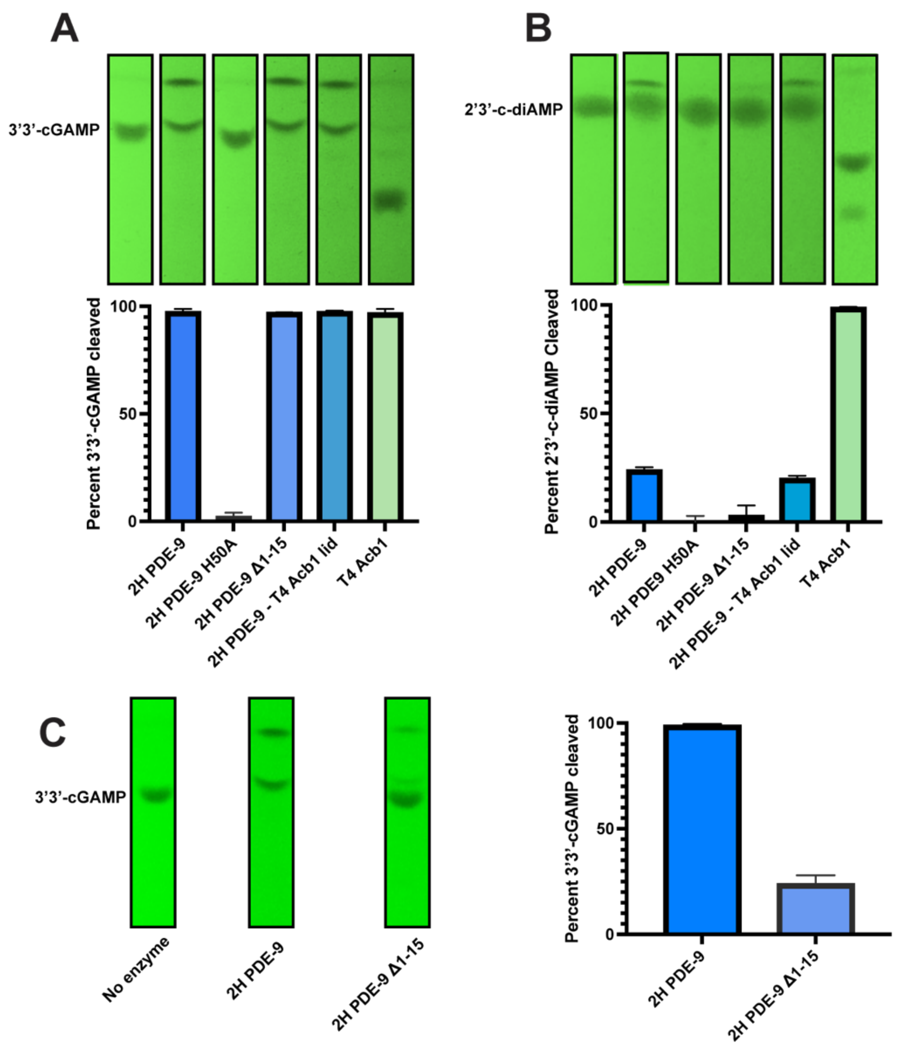


**Supplementary Fig. 12. Representative TLCs for quantification of substrate degradation in vitro. A)** Result of 1 hour reaction between 3'3'-cGAMP and PDE. Left to right TLCs correspond to the bar graph below: No enzyme, 2H PDE-9 H50A, 2H PDE-9 Δ1-15, 2H PDE-9 Δ1-15 + T4 Acb1 lid, T4 Acb1. TLCs are representative of technical triplicate. Related to Fig. 3H. **B)** Result of 1 hour reaction between 2'3'-c-diAMP and PDE. Left to right TLCs correspond to the bar graph below: No enzyme, 2H PDE-9 H50A, 2H PDE-9 Δ1-15, 2H PDE-9 Δ1-15 + T4 Acb1 lid, T4 Acb1. TLCs are representative of technical triplicate. Related to Fig. 3G-H. **C)** Left: Degradation of 3'3'-cGAMP by no enzyme, 2H PDE-9, and 2H PDE-9 Δ1-15. Right: Quantification of 3'3'-cGAMP cleaved by 2H PDE-9 and 2H PDE-9 Δ1-15.
