## Supplementary Table 3 for "Divergent viral phosphodiesterases for immune signaling evasion"

**Supplementary Table 3. X-ray data collection and refinement statistics**

| PDB ID | | 2H PDE-9 wildtype<br>9Q2G |
|---|---|---|
| **Data collection**[a,b] | | |
| Space group | | $P4_3$ |
| Cell Dimensions | | |
| | *a, b, c* (Å) | 64.968, 64.968, 50.426 |
| | α, β, γ (°) | 90, 90, 90 |
| Resolution (Å) | | 64.97-2.00 (2.05-2.00) |
| $R_{merge}$ (%) | | 6.2 (14.5) |
| $R_{pim}$ (%) | | 2.8 (9.0) |
| $CC_{1/2}$ (%) | | 99.8 (97.0) |
| $\langle I/\sigma I \rangle$ | | 18.4(7.5) |
| Completeness (%) | | 95.1 (73.8) |
| Redundancy | | 5.7 (3.2) |
| Wilson *B*-factor (Å$^2$) | | 15.18 |
| **Refinement and Validation** | | |
| Resolution (Å) | | 64.97-2.00 (2.05-2.00) |
| Unique Reflections | | 13,622 (770) |
| Number of atoms | | |
| | Protein | 1,326 |
| | Ligand | 18 |
| $R_{work}/R_{free}$ (%) | | 14.77/19.60 |
| R.m.s. deviations | | |
| | Bond lengths (Å) | 0.006 |
| | Bond angles (˚) | 0.793 |
| Poor rotamers (%) | | 0.00 |
| Ramachandran plot | | |
| | Favored (%) | 99.38 |
| | Allowed (%) | 0.62 |
| | Disallowed (%) | 0 |
| Average *B*-factor (Å$^2$) | | 19.6 |

[a]For each structure reported, data were derived from a single crystal.
[b]Numbers in parentheses correspond to the highest resolution shell
