## Supplementary Table 4 for "Divergent viral phosphodiesterases for immune signaling evasion"

**Supplementary Table 4. X-ray data collection and refinement statistics**

| PDB ID | | 2H PDE-9 wildtype<br>9Q2Y |
|---|---|---|
| **Data collection**[a,b] | | |
| Space group | | $P2_12_12_1$ |
| Cell Dimensions | | |
| | *a, b, c* (Å) | 72.513, 74.8, 76.727 |
| | α, β, γ (°) | 90, 90, 90 |
| Resolution (Å) | | 43.08-2.03 (2.08-2.03) |
| $R_{merge}$ (%) | | 16.6 (147.2) |
| $R_{pim}$ (%) | | 5.9 (52.0) |
| $CC_{1/2}$ (%) | | 99.7 (38.5) |
| <I/σI> | | 12.0(1.5) |
| Completeness (%) | | 100.0 (100.0) |
| Redundancy | | 8.8 (8.8) |
| Wilson *B*-factor (Å$^2$) | | 27.93 |
| **Refinement and Validation** | | |
| Resolution (Å) | | 43.08-2.03 (2.08-2.03) |
| Unique Reflections | | 27,651 (2023) |
| Number of atoms | | |
| | Protein | 2,573 |
| | Ligand | 11 |
| $R_{work}/R_{free}$ (%) | | 22.58/24.49 |
| R.m.s. deviations | | |
| | Bond lengths (Å) | 0.007 |
| | Bond angles (˚) | 0.847 |
| Poor rotamers (%) | | 0.32 |
| Ramachandran plot | | |
| | Favored (%) | 96.20 |
| | Allowed (%) | 3.48 |
| | Disallowed (%) | 0 |
| Average *B*-factor (Å$^2$) | | 38.3 |

[a]For each structure reported, data were derived from a single crystal.
[b]Numbers in parentheses correspond to the highest resolution shell
